# Barley variety Quest exhibits broad-spectrum, moderate quantitative resistance against diverse *Xanthomonas translucens* pathovar translucens

**DOI:** 10.1101/2024.06.09.598138

**Authors:** Nathaniel Heiden, Everett Burns, Jules Butchacas, Veronica Roman-Reyna, Kelly Mikhail, Jonathan M. Jacobs

**Affiliations:** Department of Plant Pathology, The Ohio State University, Columbus, Ohio, USA; Infectious Diseases Institute, The Ohio State University, Columbus, Ohio, USA; Department of Plant Pathology and Environmental Microbiology, Pennsylvania State University, University Park, Pennsylvania, USA; Department of Horticulture and Crop Science, The Ohio State University, Ohio, USA

## Abstract

Barley (*Hordeum vulgare* subspp. *vulgare*) is the fourth most widely produced cereal today and is valuable for both animal and human consumption. Bacterial leaf streak (BLS) of barley is caused primarily by *X. translucens* pv. translucens (Xtt) and can lead to significant yield losses. There are currently no available control methods for BLS pathogens, but there are ongoing efforts to identify and characterize sources of resistance to Xtt in available barley germplasm. These screening projects require field trials which are time-consuming and challenged due to variables introduced by environmental and weather conditions, field conditions and other organisms that may be present. Reliable greenhouse phenotyping techniques are needed to accelerate screening of barley germplasm against more Xtt strains. In this study, we established a rapid greenhouse spray inoculation protocol for pathogen-plant phenotyping. This provides a framework for rapid germplasm screening before time-consuming and limited field nursery trials. Our method confirmed the moderate quantitative resistance phenotype discovered in field trials for the cultivars Quest and Tradition against the Xtt strain CIX95. One week old seedlings of the previously characterized resistant line PI329000 were not resistant relative to the susceptible check line MW14-5371-013. The cultivar Quest then demonstrated moderate quantitative resistance against a diverse panel of Xtt strains. Xtt subgroup did not influence BLS outcomes on tested barley lines. We found evidence to support the hypothesis that the virulence of modern strains is increasing, though this does not improve their ability to cause disease on Quest.

## Introduction

Barley (*Hordeum vulgare* subspp. *vulgare*) is the fourth most widely produced cereal today and grows robustly in diverse environments and under stressful conditions [1,2]. Barley is used for animal and human consumption with significant health benefits for humans according to the USA Food and Drug Administration [2]. Barley has a history of cultivation that stretches for 10,000 years [3]. This has resulted in a high diversity of barley germplasm available to us across temporal and geographic scales [4]. This diversity enables breeders to find sources of resistance to both abiotic and biotic stressors that can be transferred into cultivated varieties (cultivars) of barley or into other important Poaceae crops.

Bacterial leaf streak (BLS) is a disease caused by *Xanthomonas* species on monocotyledonous plants and is characterized by watersoaked and later chlorotic interveinal streaks. Although there is limited reporting, BLS can lead to significant yield losses [5–12]. Limited research suggests that BLS pathogens are seedborne but can spread locally in the form of exudates via wind and water [6,7,9–11,13].

Barley BLS is caused by *X. translucens*, which is a diverse species consisting of multiple pathovars. *X. translucens* can be divided into three main clades on the basis of average nucleotide identity and DNA-DNA hybridization [14,15]. Clade Xt-I includes the two pathovars which are typically found infecting barley: translucens and undulosa [14]. Barley is threatened primarily by the pathovar translucens (Xtt) [11,16–19]. The pathovar undulosa mainly infects wheat along with a wide range of alternative hosts but has also been documented to occasionally cause BLS in barley [20,21]. Xtt is the main concern in barley production and restricted to members of the *Hordeum* genus [11,22].

BLS is a sporadic disease [17]. The incidence of BLS is associated with high temperatures and humidity with these conditions favoring the replication and spread of Xtt [12,13,17,23]. In the recent decade the threat and prevalence of BLS in barley has increased especially in the Northern Great Plains of the USA [17]. There is still debate about why BLS is an increasing problem. One explanation is that climate change has led to higher temperatures and humidity in barley-growing regions, which could favor Xtt. Changes in climate can also impact Xtt transmission as *X. translucens* has previously been isolated from precipitation [24,25]. Changes in production practices such as reduced tilling may play a role [4]. Yet another hypothesis is that Xtt strains have evolved to have increased virulence and transmissibility [26].

There are limited options to combat bacterial pathogens. Copper has been used; however *Xanthomonas* species are capable of acquiring resistance to copper [27–32]. Therefore, this is unlikely to be a sustainable technique. In line with this challenge to manage *Xanthomonas* species in general, there are currently no available control methods for BLS pathogens [17]. The gold standard for prevention of losses due to *Xanthomonas* BLS pathogens could be genetic resistance [33].

In response to this threat, there are ongoing efforts to identify and characterize sources of resistance to Xtt in available barley germplasm. Wild *Hordeum* species are also potential sources of resistant germplasm [24]. The barley variety Morex was classified as moderately resistant and several quantitative trait loci responsible for this resistance were identified in 1994 [34]. Morex is a six-rowed malting barley which, according to the USDA GrainGenes database is the “American malting industry standard”. The Morex genome is the model genome for barley as well [35]. However, it is unknown how Morex interacts with the current Xtt population. Recent analysis has separated Xtt into three separate genomic groups [36,37], but little is known about phenotypic differences among these groups. It is important therefore to examine Morex and other identified resistant barley in the context of the current Xtt population.

A recent study in Minnesota by Ritzinger et al. (2022) found multiple sources of resistance. A follow-up genome-wide association analysis found eight quantitative trait loci for BLS resistance [38]. Field trials are likely an optimal technique to screen for resistance that is truest to natural conditions. However, field trials are time consuming in nature and are challenged due to variables introduced by environmental and weather conditions, field conditions and other organisms that may be present. This means that decisions often have to be made based off of a single field season if usable data is generated. Due to these challenges, one has to limit the treatments on the pathogen side when screening a large set of germplasm.

In Ritzinger et al. (2022) some promising sources of resistance were found against the single strain CIX95, however, it is unknown if these sources of resistance are effective against the broader Xtt population. Reliable greenhouse phenotyping techniques are needed to accelerate screening of barley germplasm against more Xtt strains [11].

In this study we established a rapid greenhouse spray inoculation protocol for pathogen-plant phenotyping. We determined which days plants should be rated with this protocol to enable the application of a standard percentage rating system. We then challenged barley varieties with known resistance phenotypes with the single Xtt strain CIX95 to mirror field studies. Finally, we challenged a resistant cultivar with a diverse panel of Xtt. This overall provides a framework for rapid germplasm prescreening before time-consuming and limited field nursey trials.

## Materials and Methods

### Plant Growth Conditions

Barley seeds were planted at approximately 3cm of depth in PROMIX potting soil (Premier Tech, 2023) in 7.63cm by 7.63cm by 7.63cm pots. Experiments were completed in a greenhouse with natural lighting and temperature maintained between 21 – 26°C or in a growth chamber with temperature maintained at 26°C and humidity at 60%. 7-day old seedlings were used for all inoculations.

### Bacterial inoculum preparation

*X. translucens* strains were struck out from −80°C freezer stock onto nutrient agar (NA, 3g/L beef extract, 5g/L peptone, 15g/L agar) plates and allowed to grow for three days at 28°C. Bacteria were re-struck onto a new plate of NA media and allowed to grow for 24 hours. A 10μL loop of bacteria from the fresh growth plate was suspended in 5mL of deionized water in a 15mL conical tube. The optical density at 600nm (OD600nm) was adjusted to 1.0.

### Spray inoculation

A marker was used to delineate a 3cm section of each leaf to be inoculated. A handheld perfume sprayer was placed into a 15mL conical tube containing bacterial inoculum or a water control. The inoculum was sprayed five times on the abaxial and axial sides of the leaf. Gloves were disinfected with 75% ethanol when finished with each treatment. The inoculated plants were covered for three days after inoculation to maintain a humid environment.

### Symptom Rating for AUDPC calculation

Symptoms were rated daily for a period of seven days past inoculation (dpi). Symptoms were classified on a scale from zero to four considering only the delineated inoculated zone on a leaf. A rating of zero referred to no symptoms. A rating of one signified the presence of discrete circular or oval water-soaking spots. A rating of two meant that the initially formed translucent lesions began to enlarge into streaks but remained discrete. A rating of three described when watersoaking lesions formed streaks without individual watersoaking spots visible. A rating of four signified the presence of chlorosis and/or necrosis. A disease progress curve was constructed in R version 4.3.1 [39] and the R package agricolae [40] was used to calculate area under the disease progress curve (AUDPC). A linear regression was run in R version 4.3.1 [39] to compare each individual timepoint to the total AUDPC values.

### Single day disease rating

Pictures were taken of leaf symptomatic area at five dpi. Percent symptomatic area was calculated with ImageJ [41]. In ImageJ the total leaf area was selected and the total pixel count in that area was calculated with the area tool. Then, only the symptomatic area was selected and the total pixel count in the symptomatic area was calculated with the area tool. A ratio of symptomatic area to total area was calculated to provide a percentage. Percentages were converted to ratings on a scale from one to nine based on Alizadeh et al. (1994).

### Statistical Analysis

All statistical analyses were conducted in R version 4.3.1 [39]. Mean separations from normally distributed datasets were determined via an ANOVA and then Tukey Honest Significant Differences test. For non-normally distributed data, model significance was tested with a Kruskal-Wallis rank sum test and mean separation for multiple comparisons was conducted using a Wilcoxon rank sum test with continuity correction.

## Results

### Development of rapid germplasm pre-screening method

Field level resistance screening of *X. translucens* on barley and other cereal hosts is typically performed by backpack spray inoculation [4]. This is designed to mimic natural *X. translucens* infection. This resistance screening is limited to one time annually pending weather or other events in a barley production zone. Therefore, we adapted this field protocol to a greenhouse setting to quantify symptoms quickly and reproducibly in barley. It was important for this approach to be applicable to controlled environment settings to allow year-round screening. Instead of a large gas-powered backpack sprayer, we developed a protocol using a handheld perfume sprayer. We used one week-old plants and focused inoculation on a single zone of each inoculated leaf to allow for rapid and reproducible ratings.

Due to practical restrictions in the aforementioned field trials, disease is usually rated at a single timepoint. It is unknown for these assays which timepoint is most predictive of the overall disease outcome. We therefore created a rapid screening technique that would enable us to assess disease on a daily basis (Fig 1A). We challenged the model barley cultivar Morex (Table 1) with Xtt strains CIX392, CIX394, CIX401, CIX43, CIX95, SLV-2, UPB458, UPB787, UPB820, UPB886 and UPB906 and the pv. undulosa strains CIX184, CIX354, CIX40, UPB513 and UPB882 (Table 2; Fig 1B). Disease progress curves generated from daily ratings were different for each strain (Fig 1B; Fig S2) which resulted in AUDPC values ranging from zero to 15 (Fig S2).

**Figure 1.**
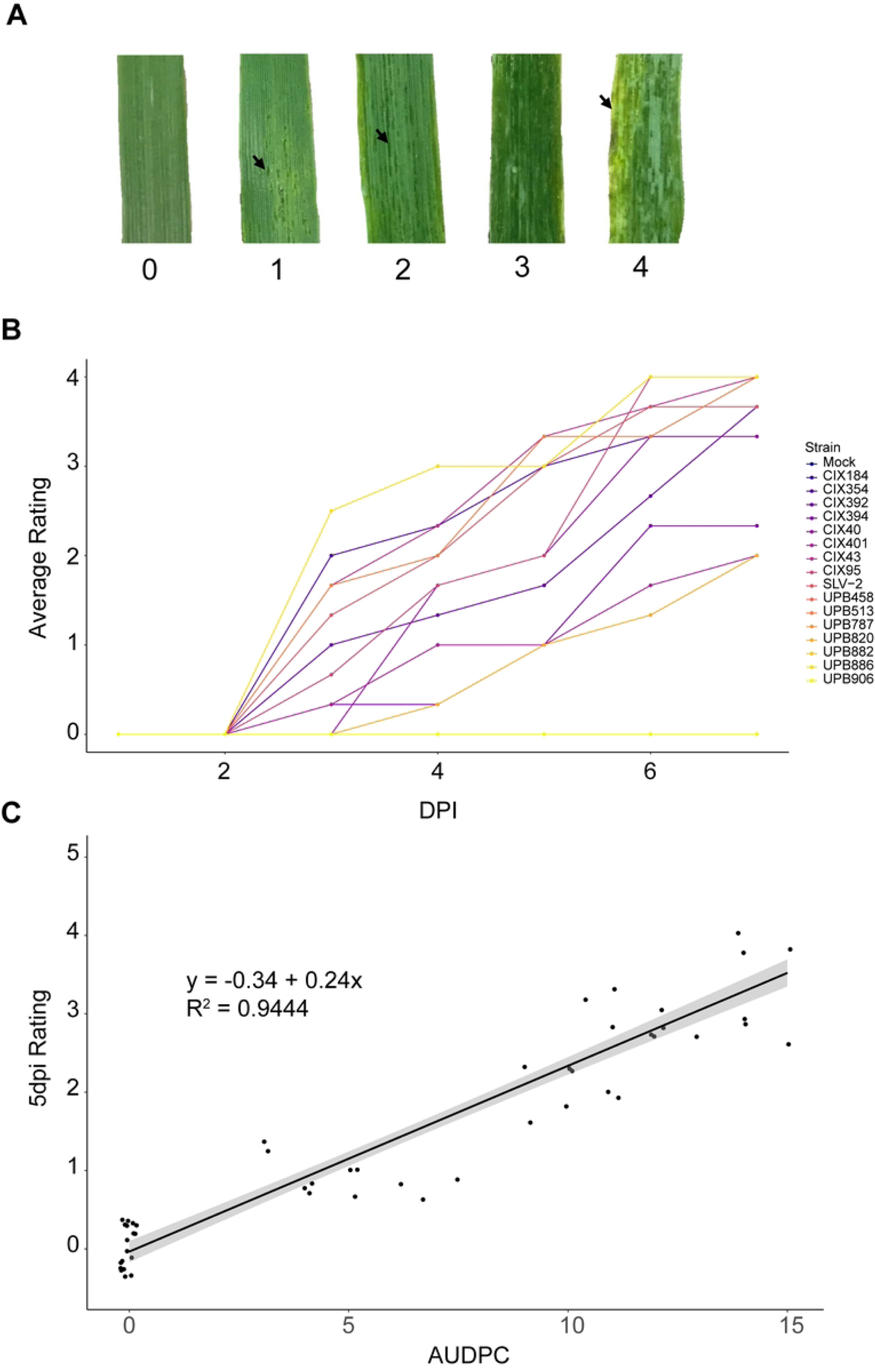
BLS rating at 5 days post inoculation describes Xtt-barley interaction. (A) Representative images of ratings used for rating system. A rating of zero referred to no symptoms. A rating of one signified the presence of discrete circular or oval water-soaking spots. A rating of two meant that the initially formed translucent lesions began to enlarge into streaks but remained discrete. A rating of three described when watersoaking lesions formed streaks without individual watersoaking spots visible. A rating of four signified the presence of chlorosis and/or necrosis. (B) Barley cultivar Morex was spray inoculated with *X. translucens* strains CIX184, CIX354, CIX392, CIX394, CIX40, CIX401, CIX43, CIX95, SLV-2, UPB458, UPB513, UPB787, UPB820, UPB882, UPB886 and UPB906 at 1.0 O.D. 600nm or with a water mock control. Average disease ratings are displayed were for each strain per day and displayed as average disease progress curves with R version 4.3.1 [39]. (C) The R package agricolae [40] was used to calculate area under the disease progress curve (AUDPC) for each replicate inoculated leaf zone. A linear regression was run in R version 4.3.1 [39] to compare each individual timepoint to the total AUDPC values.

**Table 1.** List of Xtt strains used in this study.

| Strain | Pathovar | Xtt subgroup | Year Isolated | Location Isolated | Publication |
| --- | --- | --- | --- | --- | --- |
| BSLB2 | translucens | K1 | 2017 | North Dakota, USA | NDSU, unpublished |
| CFBP2054 (NCPPB973) | translucens | K0 | 1933 | Minnesota, USA | Bragard et al. 1995 |
| CIX257 | translucens | K1 | 2017 | Minnesota, USA | Ledman, unpublished |
| CIX261 | translucens | K2 | 2016 | Minnesota, USA | Ledman et al. 2021 |
| CIX392 (B1.6) | translucens | K0 | 2015 | Minnesota, USA | Curland et al. 2020 |
| CIX394 (B1.8) | translucens | K0 | 2015 | Minnesota, USA | Curland et al. 2020 |
| CIX401 (B1.15) | translucens | unclassified | 2015 | Minnesota, USA | Curland et al. 2020 |
| CIX43 | translucens | K0 | 2009 | Minnesota, USA | Curland et al. 2018 |
| CIX72 | translucens | K2 | 2006 | Minnesota, USA | Curland et al. 2018 |
| CIX81 | translucens | K0 | 2011 | North Dakota, USA | Curland et al. 2018 |
| CIX90 | translucens | K2 | 2011 | Minnesota, USA | Curland et al. 2018 |
| CIX95 | translucens | K0 | 2011 | Minnesota, USA | Curland et al. 2018 |
| CIX96 | translucens | K0 | 2011 | North Dakota, USA | Curland et al. 2018 |
| SLV-2 (CIX75) | translucens | K1 | 2010 | Colorado, USA | Langlois et al. 2017 |
| UPB458 | translucens | K1 | 1970 | India | Bragard et al. 1997 |
| UPB545 | translucens | K1 | 1987 | Batan, Mexico | Bragard et al. 1995 |
| UPB787 | translucens | K1 | 1990 | Jica, Paraguay | Bragard et al. 1997 |
| UPB820 | translucens | K2 | 1990 | Arak, Iran | Bragard et al. 1997 |
| UPB886 | translucens | K1 | 1990 | Arak, Iran | Bragard et al. 1997 |
| UPB906 | translucens | K2 | 1990 | Kerman, Iran | Bragard et al. 1997 |
| XtKm34 | translucens | K2 | 2015 | Kerman, Iran | Khojasteh et al. 2019 |
| XtKm7 | translucens | K0 | 2014 | Kerman, Iran | Khojasteh et al. 2019 |
| XtKm8 | translucens | K2 | 2014 | Kerman, Iran | Khojasteh et al. 2019 |
| CIX184 | undulosa | N/A | 2017 | Minnesota, USA | Ledman et al. 2021 |
| CIX354 | undulosa | N/A | 2015 | Minnesota, USA | Roman-Reyna et al. 2023 |
| CIX40 | undulosa | N/A | 2010 | Minnesota, USA | Curland et al. 2020 |
| UPB513 | undulosa | N/A | 1987 | Mexico | Bragard et al. 1995 |
| UPB882 | undulosa | N/A | 1991 | Yemen | Bragard et al. 1995 |

Strains used in this study were classified as K0, K1 or K2 using the PCR assay described in
Ebeling-Koenig et al. Available metadata regarding the state and/or country was included as
available.

**Table 2.**
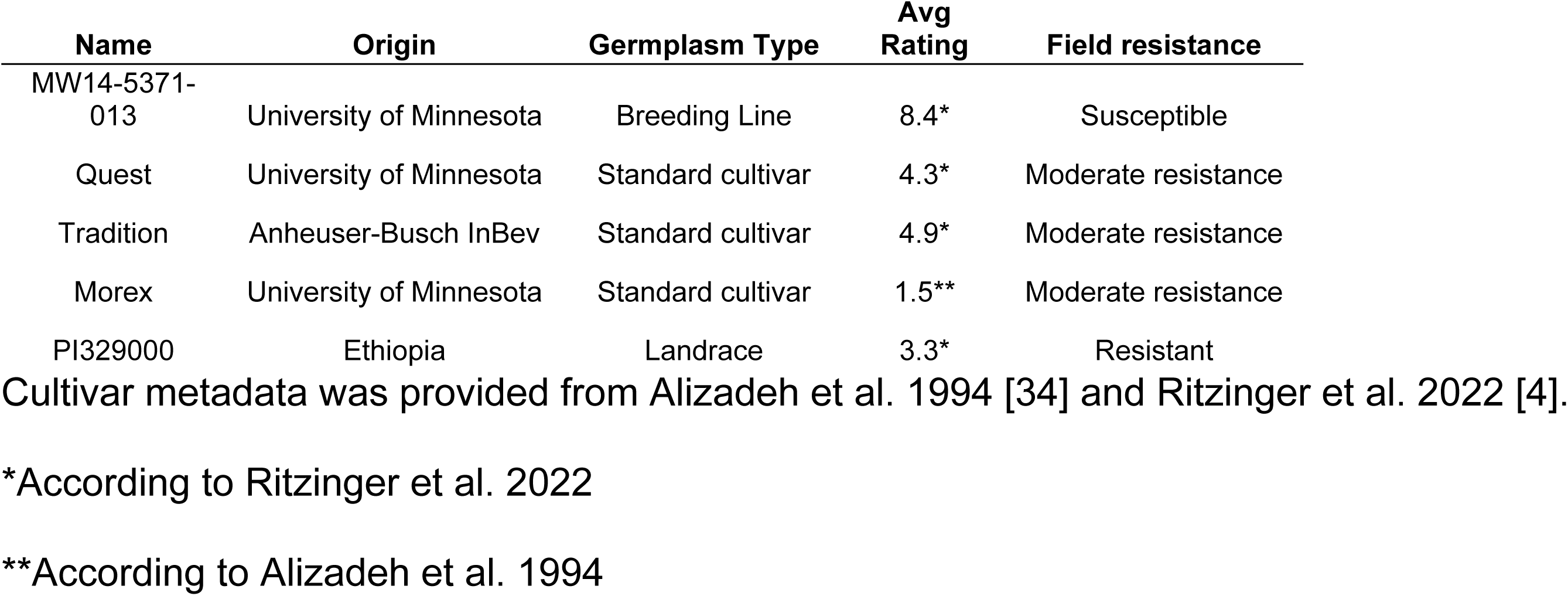
List of barley lines used in this study.

We comprehensively examined symptoms every day for seven days, which might be impractical for large scale trials with breeding populations and multiple strain types as Xtt is globally diverse [36]. We hypothesized that a single time point would correlate closely with the AUDPC. To test this hypothesis, we ran linear regressions of ratings at each time point against AUDPC for that leaf. We found a significant correlation for all days three through seven (p-value < 0.01), which is when disease symptoms were present (Fig S3). AUDPC correlated with ratings at three days had an R^2^ value of 0.7241, four days had a R^2^ value of 0.9240, five days had an R^2^ value of 0.9444, six days had an R^2^ value of 0.9372 and seven days had an R^2^ value of 0.9270 (Fig S3). Ratings at five days had the highest correlation with AUDPC according to R2 value, and so we chose five days as the timepoint to assess virulence (Fig 1C; Fig S3).

### Comparing pre-screening method on field-tested cultivars

Resistance is not available in most commercial barley production. Few barley genotypes are partially resistant according to field data, including cultivars Quest and Morex and land race PI329000 [4,34]. We therefore compared these barley lines against the line MW14-5371-013, which served as a susceptible check [4]. We inoculated these cultivars with Xtt strain CIX95 to compare to previously published field inoculations carried out with this strain [4,20]. As ratings could be completed in a single day, we used a more precise measurement of percent symptomatic area, which was then converted into the standard ratings scale used in field experiments [4,34].

Disease across replicates was greater in the first compared the second replicate, but several main trends remained consistent. The susceptible check line MW14-5371-013 was, as expected, susceptible across both replicates with an average rating of 6.67 and 2.67 (SE= 0.21, 0.14; Fig 2, Table 3). Quest was relatively resistant in both replicates with average ratings of 4.33 and 1.17 (SE = 0.17, 0.07; Figure 2, Table 3). The resistant check line PI329000 was equally as susceptible as MW14-5371-013 in both replicates (Fig 2) and averaged ratings of 6.89 and 3.5 (SE = 0.14, 0.25; Table 3). Morex has previously been reported resistant to strains not included in this study [34]. However, Morex was not resistant in our assay (Figure 2) for replicate 1 (average score = 6.00, SE = 0.5; S1 Table) or replicate 2 (average score = 1.5, SE = 0.14; Table 3). As the contemporary variety Quest was resistant in our first replicate, we included the cultivar Tradition, which is another modern barley variety with intermediate field resistance, in our second replicate. Tradition was significantly resistant against CIX95 compared to MW14-5371-013 with an average rating of 1.38 (SE = 0.06) (Figure 2, Table 3).

**Figure 2.**
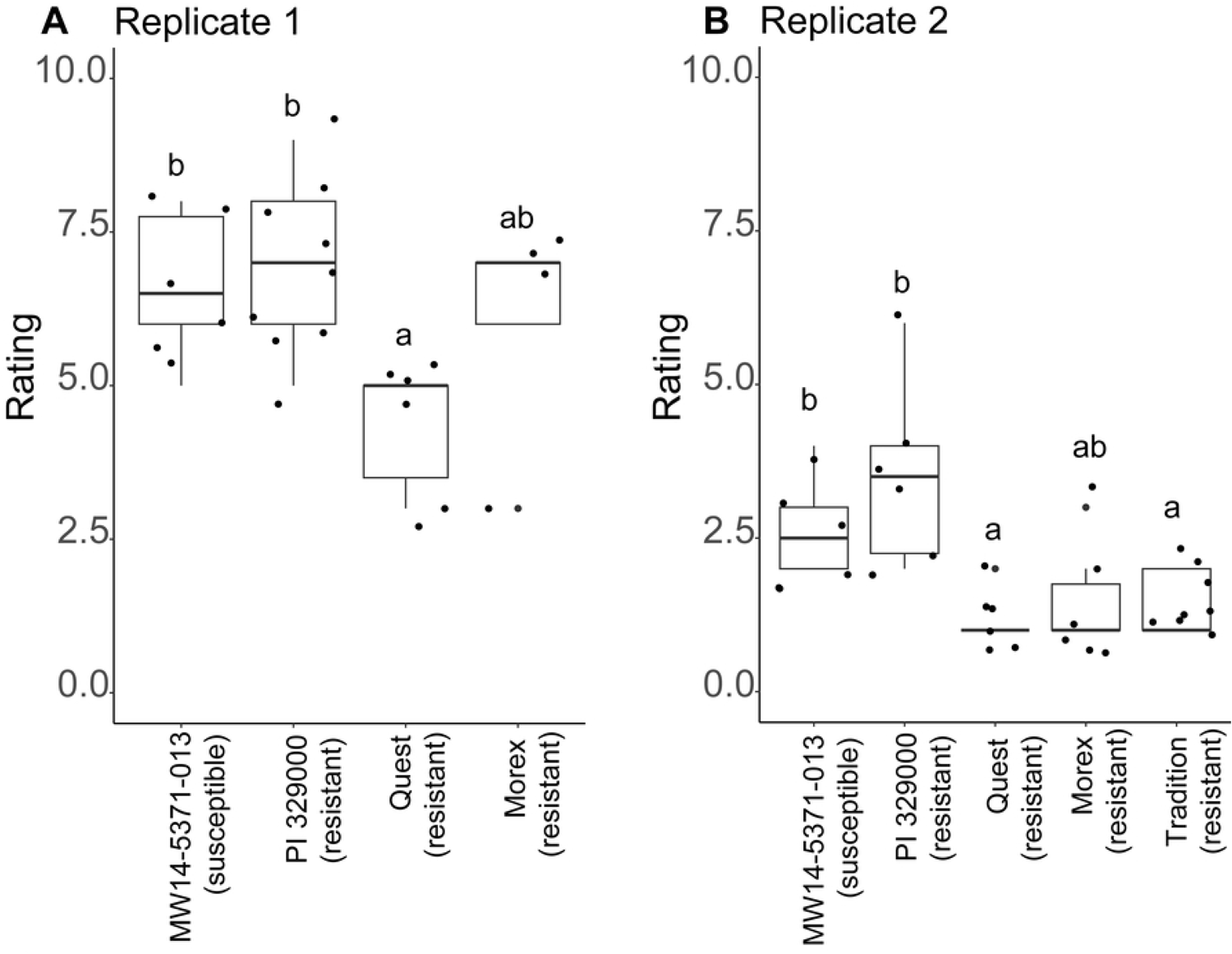
Field resistance exhibited by barley cultivar Quest is reflected in spray inoculation. Barley cultivars MW14-5371-013, PI329000, Quest and Morex were spray inoculated with Xtt strain CIX95 at 1.0 O.D.600nm and pictures of symptoms were taken at 5dpi in two separate experimental replicates. The cultivar Tradition was added to the second replicate. ImageJ was used to calculate percent symptomatic area and percent symptomatic area values were converted to ratings used in field trials [4,34]. Letters represent significant differences (p < 0.05) between groups based on pairwise comparisons using a Wilcoxon rank sum test with continuity correction in R version 4.3.1 [39].

**Table 3.**
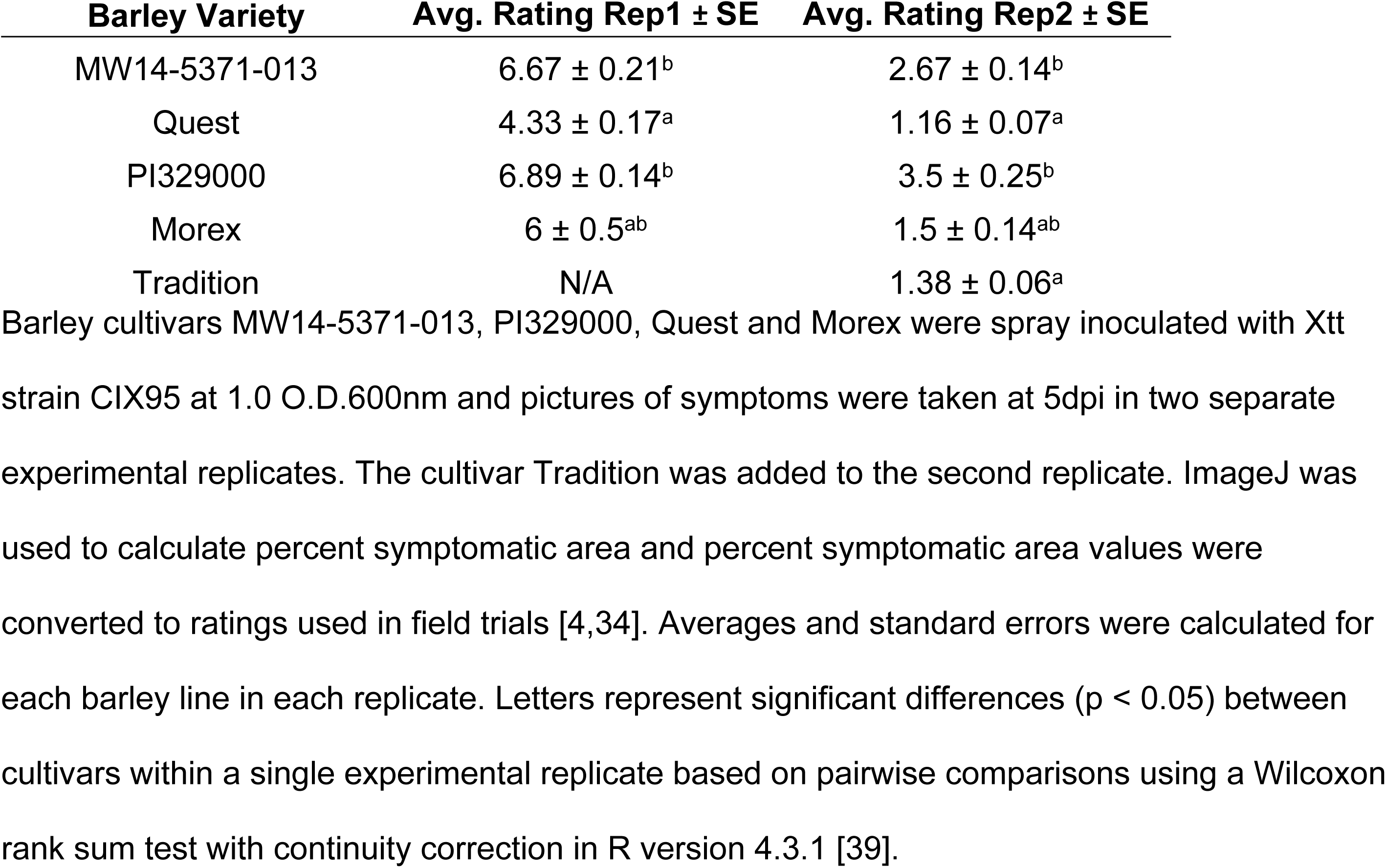
Bacterial leaf streak ratings by barley variety.

| Barley Variety | Avg. Rating Rep1 ± SE | Avg. Rating Rep2 ± SE |
| --- | --- | --- |
| MW14-5371-013 | $6.67 \pm 0.21^b$ | $2.67 \pm 0.14^b$ |
| Quest | $4.33 \pm 0.17^a$ | $1.16 \pm 0.07^a$ |
| PI329000 | $6.89 \pm 0.14^b$ | $3.5 \pm 0.25^b$ |
| Morex | $6 \pm 0.5^{ab}$ | $1.5 \pm 0.14^{ab}$ |
| Tradition | N/A | $1.38 \pm 0.06^a$ |

Barley cultivars MW14-5371-013, PI329000, Quest and Morex were spray inoculated with Xtt strain CIX95 at 1.0 O.D.600nm and pictures of symptoms were taken at 5dpi in two separate experimental replicates. The cultivar Tradition was added to the second replicate. ImageJ was used to calculate percent symptomatic area and percent symptomatic area values were converted to ratings used in field trials [4,34]. Averages and standard errors were calculated for each barley line in each replicate. Letters represent significant differences ( $p < 0.05$ ) between cultivars within a single experimental replicate based on pairwise comparisons using a Wilcoxon rank sum test with continuity correction in R version 4.3.1 [39].

### Pre-screening against broad Xtt diversity, time of isolation and geographic distribution

Data from the field [4] and our greenhouse assay demonstrate that Quest is partially resistant to infection with the Xtt strain CIX95. However, Xtt strains are diverse and include three different subgroups: K0, K1 and K2 [36,37]. We hypothesized that Quest is resistant to Xtt strains from all three subgroups. To test this hypothesis, we inoculated both Quest and the susceptible MW14-5371-013 with Xtt strains from all three groups. For group K0 we included the current strains CIX95 and CIX96 isolated in 2011 and the model strain CFBP2054 which was isolated in 1933, all from the USA. We also included the strain XtKm7 which is a modern strain isolated in Iran in 2014. In group K1 we used the modern strains BLSB2 and CIX257, isolated in the USA in 2017 and the historical strains UPB458 and UPB545. UPB458 was isolated in India in 1970 and UPB545 was isolated in Mexico in 1987. For K2 we used the modern strains CIX72, CIX90 and CIX261 which were isolated from the USA in 2006, 2011 and 2011, respectively. We also included the Iranian strains XtKm8 and XtKm34 which were isolated in 2014 and 2015, respectively.

In general, Quest was resistant to Xtt strains compared to MW14-5371-013 (Fig 3A). Quest had an average rating of 2.80 (SE = 0.04) compared to 5.78 (SE = 0.08) for MW14-5371-013 (S1 Table). With the exception of CIX96, UPB458 and CIX72, all strains had a greater average disease rating on MW14-5371-013 than on Quest (Fig 3A, Table S2). CIX96 had an average rating of 1.00 (SE = 0) on MW14-5371-013 but was more virulent on Quest with an average rating of 3.67 (SE = 0.19; Fig 3A, Table S2). UPB458 and CIX72 caused BLS symptoms on neither MW14-5371-013 nor Quest (Fig 3A) and so had average ratings of 1.00 (Table S2).

**Figure 3.**
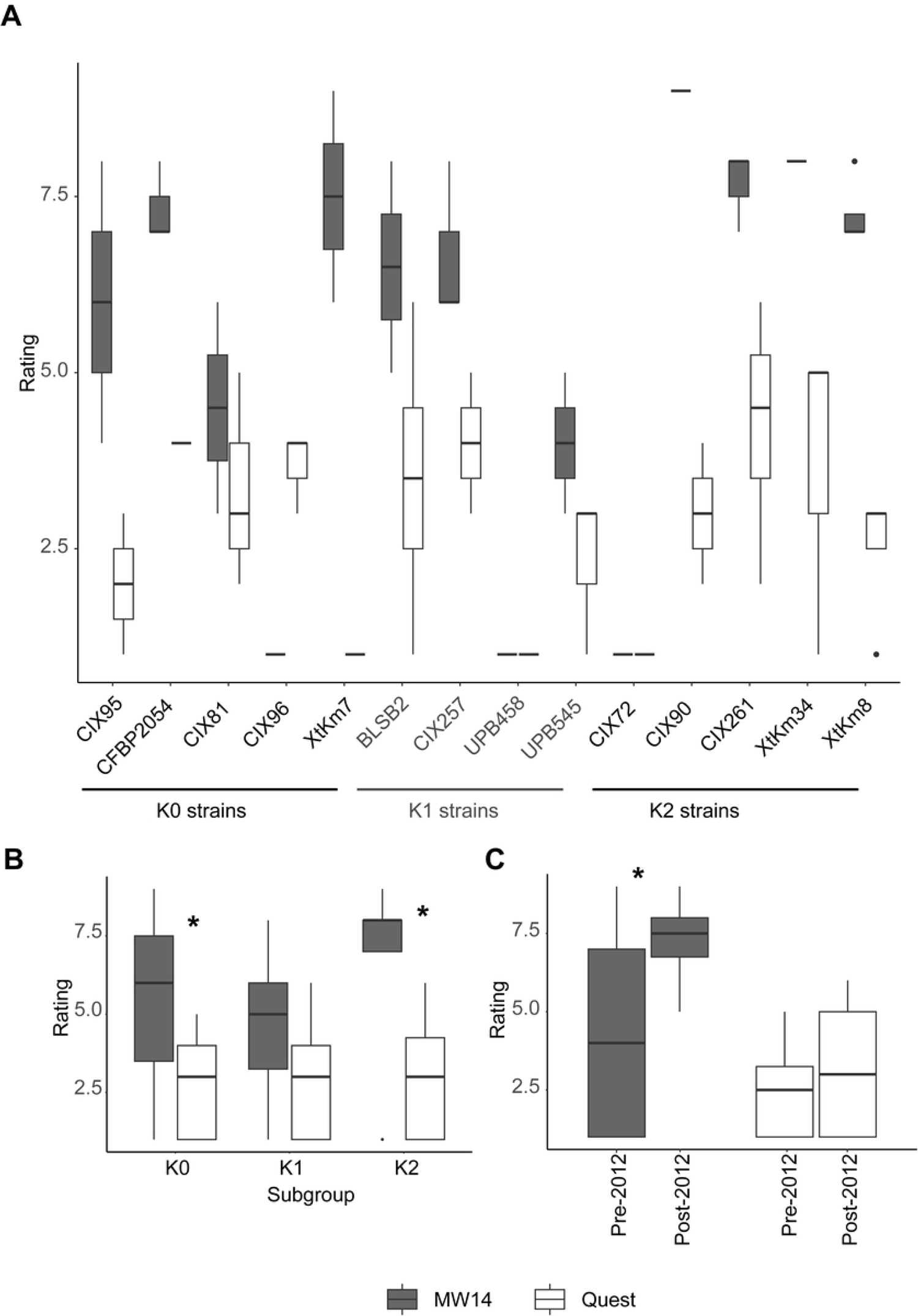
Quest resistance is broad-spectrum against diverse Xtt strains. Bolded strains are from Asia (Iran or India) remaining strains are from the U.S.A. Barley cultivars MW14-5371-013 and Quest were spray inoculated at 1.0 O.D.600nm with 14 Xtt strains: CFBP2054 (n = 3 MW14-5371-013, n = 1 Quest), CIX81 (n = 2 MW14-5371-013, n = 3 Quest), CIX96 (n = 2 MW14-5371-013, n = 3 Quest), CIX95 (n = 2 MW14-5371-013, n = 2 Quest), XtKm7 (n = MW14-5371-013, n = 3 Quest), BLSB2 (n = 2 MW14-5371-013, n = 4 Quest), CIX257 (n = 3 MW14-5371-013, n = 3 Quest), UPB458 (n = 2 MW14-5371-013, n = 3 Quest), UPB545 (n = 3 MW14-5371-013, n = 3 Quest), CIX72 (n = 1 MW14-5371-013, n = 3 Quest), CIX90 (n = 1 MW14-5371-013, n = 2 Quest), CIX261 (n = 3 MW14-5371-013, n = 4 Quest), XtKm34 (n = 2 MW14-5371-013, n = 3 Quest), XtKm8 (n = 4 MW14-5371-013, n = 4 Quest). Symptoms were assessed at 5dpi. ImageJ was used to calculate percent symptomatic area and percent symptomatic area values were converted to ratings used in field trials [4,34]. (A) Individual strains are shown on both MW14-5371-013 and Quest. (B) Strains were categorized into subgroups according to Heiden et al. (2023) with diagnostic primers [37]. Strains UPB458 and CIX72 did not have ratings above 1 on either MW14-5371-013 or Quest and were removed from the within subgroup comparisons. All strains were included in analyses described in other panels of this figure. (C) Strains were separated into groups depending on whether they were isolated before (Pre-2012) or after (Post-2012) the year 2012. An asterisk (*) signifies a significant difference (p < 0.05) based on pairwise comparisons using a Wilcoxon rank sum test with continuity correction in R version 4.3.1 [39].

We observed previously that Quest was resistant to K0 subgroup strain CIX95. Our results also demonstrate that Quest had lower disease on average compared to MW14-5371-013. According to Kruskal-Wall rank sum tests, subgroups did not have a significant impact on BLS rating on MW14-5371-013 (Kurkal-Wallis chi-squared = 5.9366, df = 2, p-value = 0.05139) and Quest (Kruskal-Wallis chi-squared = 0.16277, df = 2, p-value = 0.9218). For within subgroup analysis the strains UPB458 and CIX72 were removed as they did not have ratings above 1 on either MW14-5371-013. There was significantly greater disease in MW14-5371-013 compared to resistant Quest within subgroup K0 (Kruskal-Wallis chi-squared = 5.6311, df = 1, p-value = 0.0176; Fig 3B), K1 (Kruskal-Wallis chi-squared = 5.9645, df = 1, p-value = 0.0146; Fig 3B) and K2 (Kruskal-Wallis chi-squared = 16.586, df = 1, p-value < 0.01; Fig 3B).

Because BLS has continued to increase in prevalence as a threat to barley production in recent years, our aim was to investigate whether this trend could be due to increased virulence of modern strains. We compared symptoms from our strains that were isolated this decade and those isolated prior (Fig 3C). Our strain panel consisted of strains CIX95, CIX95, CFBP2054, UPB458, UPB545, CIX72, CIX90, CIX261 isolated prior to 2012 and strains XtKm7, BLSB2, CIX257, XtKm8 and XtKm34 isolated after 2012. On average across both MW14-5371-013 and Quest, leaves infected with strains isolated prior to 2012 had a rating of 3.25 (SE = 0.07; S1 Table) and the strains isolated after 2012 had a rating of 4.95 (SE = 0.07; S1 Table). This difference was indeed significant (Kruskal-Wallis chi-squared = 8.0466, df = 1, p-value = 0.004559), but after visual inspection (Fig 3C) we hypothesized that this was mostly explained by virulence differences between modern and historical strains on the susceptible barley variety MW14-5371-013. In support of our hypothesis, on MW14-5371-013 alone, strains isolated after 2012 produced significantly greater BLS symptoms than those isolated before 2012 (Kruskal-Wallis chi-squared = 8.1668, df = 1, p-value = 0.004266; Fig 3C, S1 Table). However, on Quest alone there was no significant difference in BLS rating between strains isolated before and after 2012 (Kruskal-Wallis chi-squared = 2.152, df = 1, p-value = 0.1424). This result suggests that the virulence of modern strains may be increasing, though this does not improve their ability to cause disease on Quest.

## Discussion

In this study we developed a rapid greenhouse pre-screening spray inoculation method using one week old plants and focused inoculation on a single zone of each leaf. We created preliminary rating system and tracked disease over time which allowed us to determine that five dpi was the best timepoint to assess Xtt-barley interactions. We then quantitatively assessed the interaction of the strain CIX95, used in field trials, against multiple Xtt strains. This trial found that cultivar Quest is resistant in our greenhouse assay as in the field setting.

Our pre-screening method can be used to provide rating values on the same scale as that used in field trials. It built on the method established initially by Alizadeh et al. [34] and focused on inoculation of a single area of a leaf and the implementation of image processing software. Field trials will continue to be necessary to confirm resistance phenotypes. However, this high-throughput method can be used to complement and inform field trials to screen for BLS resistance against more diverse sets of Xtt.

Our research [36,37] and that of other groups [15,18–21,42] has shown that Xtt is diverse and consists of three different clades which we refer to as K0, K1 and K2. In this trial we confirmed field results [4] showing that barley cultivar Quest is resistant against a single strain of Xtt and then challenged Quest with a broad panel of Xtt. Our results demonstrated that Quest is resistant to Xtt from all three groups of Xtt. This data needs to be confirmed through multiple seasons of growth trials, however as Quest is an available cultivar growers can consider deploying Quest in lieu of other more susceptible varieties. Quest is largely used in the USA, but farmers in other countries could consider its use, as all three subgroups of Xtt are broadly distributed and Quest provided resistance against international strains such as XtKm8, XtKm34 and XtKm7 from Iran. Again, field trials against native populations of Xtt specific to a region should be carried out to confirm this.

Quest does not exclude Xtt nor prevent BLS completely, making this an example of quantitative or incomplete resistance. Though this type of resistance can allow yield losses to continue to occur, it is often a more sustainable form of resistance than complete resistance. This is because quantitative resistance does not provide as much evolutionary pressure on a pathogen, such as Xtt. As an example, this type of pressure was placed on *X. oryzae* populations when rice resistance genes were deployed, which induced major shifts in the *X. oryzae* population to favor strains that could overcome this resistance and therefore survive [43]. The use of varieties such as Quest pose little threat to drive increased virulence of the Xtt population. Full resistance to *X. translucens* by barley has never been documented. Therefore, any discovery of asymptomatic strains is of interest. Further research into the potentially incompatible interactions of Km7 on Quest, CIX96 on MW14-5371-013 and UPB458 and CIX72 on both Quest and MW14-5371-013 will may provide insight on mechanisms employed by barley lines which prevent BLS.

A benefit of this greenhouse phenotyping method is that we can examine the susceptibility of barley in different growth stages. In our study we use one week old plants. We therefore saw that Quest resistance is effective in a one-week-old seedling. The landrace PI329000 is resistant at an older life stage in the field, but this resistance was not present in one week old seedlings. We do not know if early-stage resistance is preferential to resistance that is only effective in more developed plants. Disease in Xtt typically presents itself at later stages of growth. This coincides with the warmer temperatures later in barley growing seasons. However, the transmission of Xtt is still not well understood. Seed transmission is thought to be an important contributor to BLS incidence [44]. If transmission occurs via seeds, then infection likely will occur at the earliest stages of barley infection. It is unknown if barley plants are infected with a latent infection in cooler temperatures that accelerates under more ideal conditions. We hypothesize that early-stage infection is a significant contributor to BLS and therefore characterized resistance should be examined at an early stage.

Researchers in Colorado isolated an Xtt strain named CO236 from an epidemic in Colorado and found that this strain caused more watersoaking on the cultivar Morex than the historical isolate UPB886 [26]. Based on this information, Gutierrez-Castillo et al. hypothesized that the *X. translucens* population may be increasing in virulence [26]. The authors, however, did not find evidence that there were differences in colonization ability or virulence factors compared to the Xtt population at large [26]. We found some evidence that current strains are more virulent than historical strains. Several of the historical strains used in our study did not cause symptoms or caused only minimal symptoms. We cannot discount the possibility that decreased virulence of historical strains is an artifact of their long-term storage and repeated passaging on artificial media.

The *X. translucens* research community has access to limited historical strains that were sporadically isolated and may not represent the Xtt population that has historically been present. Large isolation efforts have been undertaken that increase our understand of Xtt populations in recent years [18,20,21,26,37,42,45–48]. As additional strains are isolated in the next few years, we will be able to compare the virulence of these strains with well-characterized and diverse strain libraries from the 2000s and 2010s that are now available. It will continue be important to screen deployed germplasm against updated Xtt panels.

## Acknowledgements

The authors would like to thank Yesenia Velez-Negron for assistance with growing barley cultivars. We thank Isaac Knowles and Michael Kelly for their assistance with growing plants. We thank Mitchell G. Ritzinger, Yoonjung Lee, Dr. Brian J. Steffenson, Rebecca D. Curland and Dr. Ruth Dill-Macky for providing barley seeds and *Xanthomonas translucens* strains for this study.

## Supporting information

**S1 Fig. Strains have different disease progress curves on the same barley cultivar.** Barley cultivar Morex was spray inoculated with *X. translucens* strains CIX184, CIX354, CIX392, CIX394, CIX40, CIX401, CIX43, CIX95, SLV-2, UPB458, UPB513, UPB787, UPB820, UPB882, UPB886 and UPB906 at 1.0 O.D. 600nm or with a water mock control. Symptoms were rated on a daily basis from one to four. A rating of one signified the presence of discrete circular or oval water-soaking spots. A rating of two meant that the initially formed translucent lesions began to enlarge into streaks but remained discrete. A rating of three described when watersoaking lesions formed streaks without individual watersoaking spots visible. A rating of four signified the presence of chlorosis and/or necrosis. The R package agricolae [40] was used to calculate area under the disease progress curve (AUDPC) for each replicate inoculated leaf zone. AUDPC values are displayed under each curve which were constructed in R version 4.3.1 [39].

**S2 Fig. Ratings at day five correlate the most closely with area under the disease progress curve.** Barley cultivar Morex was spray inoculated with *X. translucens* strains CIX184, CIX354, CIX392, CIX394, CIX40, CIX401, CIX43, CIX95, SLV-2, UPB458, UPB513, UPB787, UPB820, UPB882, UPB886 and UPB906 at 1.0 O.D. 600nm or with a water mock control. Symptoms were rated on a daily basis from one to four. A rating of one signified the presence of discrete circular or oval water-soaking spots. A rating of two meant that the initially formed translucent lesions began to enlarge into streaks but remained discrete. A rating of three described when watersoaking lesions formed streaks without individual watersoaking spots visible. A rating of four signified the presence of chlorosis and/or necrosis. The R package agricolae [40] was used to calculate area under the disease progress curve (AUDPC) for each replicate inoculated leaf zone. Linear regressions were run for ratings at each day against the total AUDPC for each replicate in R version 4.3.1 [39].

**S1 Table. Bacterial leaf streak ratings by strain groupings and barley variety**

**S2. Table. Bacterial leaf streak ratings by strain and barley variety**

